# usiGrabber: Automating the curation of proteomics spectra data at scale, making large datasets ready for use in machine learning systems

**DOI:** 10.64898/2026.03.15.711873

**Authors:** Georg Auge, Matthis Clausen, Konstantin Ketterer, Jacob Schaefer, Nils Schmitt, Tom Altenburg, Yannick Hartmaring, Hendrik Raetz, Christoph N. Schlaffner, Bernhard Y. Renard

## Abstract

**Motivation:** An unprecedented amount of mass spectrometry-based proteomics data is publicly available through repositories such as the PRoteomics IDEntifications Database (PRIDE), and the field is increasingly leveraging machine learning approaches. However, the available data is not ready to be reused in a scalable way beyond the original acquisition purpose. Existing machine learning models commonly rely on a few manually curated datasets that require deep domain expertise and tedious technical work to construct. Importantly, these datasets have not been updated in recent years, so that newly published data remains inaccessible. We present usiGrabber, a scalable framework for assembling large proteomic datasets. usiGrabber is designed around portability and extensibility. It extracts spectra identification data from mzIdentML files, stores additional project-level metadata retrieved through the PRIDE API, indexes raw spectra using Universal Spectrum Identifiers (USIs), and offers download utilities to retrieve spectra data at scale.

**Results:** Within 49 hours, we parsed over 800 million peptide spectrum matches and corresponding USIs from over 1,200 projects. As a proof of concept, we used usiGrabber to construct a phosphorylation-specific training dataset of nearly 11 million spectra in under two days and used it to retrain a binary phosphorylation classifier based on the AHLF model architecture. With a balanced accuracy of 0.78, our model achieves comparable performance to the original model on an independent test set, showing that automated data extraction is an alternative to manual curation of static datasets.

**Availability:** All code is available at https://github.com/usiGrabber/usiGrabber; the data is available at https://zenodo.org/records/18853258.

## Introduction

The proteomics community has produced an unprecedented amount of publicly accessible mass spectrometry-based data, and the rate of new submissions continues to accelerate (15). In the PRoteomics IDEntifications Database (PRIDE) alone, 864 TB of data were submitted in 2025.^1^ In the same year, PubMed indexed 1,094 publications that match “machine learning proteomics”, a number that has been growing exponentially with an average of 32 % year-over-year for the last five years^2^. Still, machine learning in proteomics remains underused compared to other fields of omics research.

In practice, much of the newly available data is not ready to be leveraged en masse for machine learning. Constructing a task-specific dataset typically requires both deep domain expertise and substantial technical effort. Currently, researchers manually select relevant spectra from accessible projects, leaving out the remaining data and projects. This workflow is outlined in Figure 1A. Even for experienced researchers, curation remains time-intensive (13) and often infeasible at scale. As a result, many recent works continue to rely either on synthetic data (17) or on a small set of well-known datasets that are several years old. For example, both Casanovo (24) and Modanovo (8), published in 2024 and 2025 respectively, tackle de novo peptide sequencing from tandem mass spectra, with Modanovo explicitly extending the approach to be post-translational modification–aware for modified peptides. Yet, both still use MassIVE-KB v1 (June 2018) (22), which is now nearly eight years old. AHLF (1), published in 2022, likewise learns directly from fragmentation spectra for tasks such as detecting phosphorylated (and cross-linked) peptides and for rescoring to improve identifications at fixed error rates, but it relies on a dataset collected in June 2017 (13).

**Figure 1.**
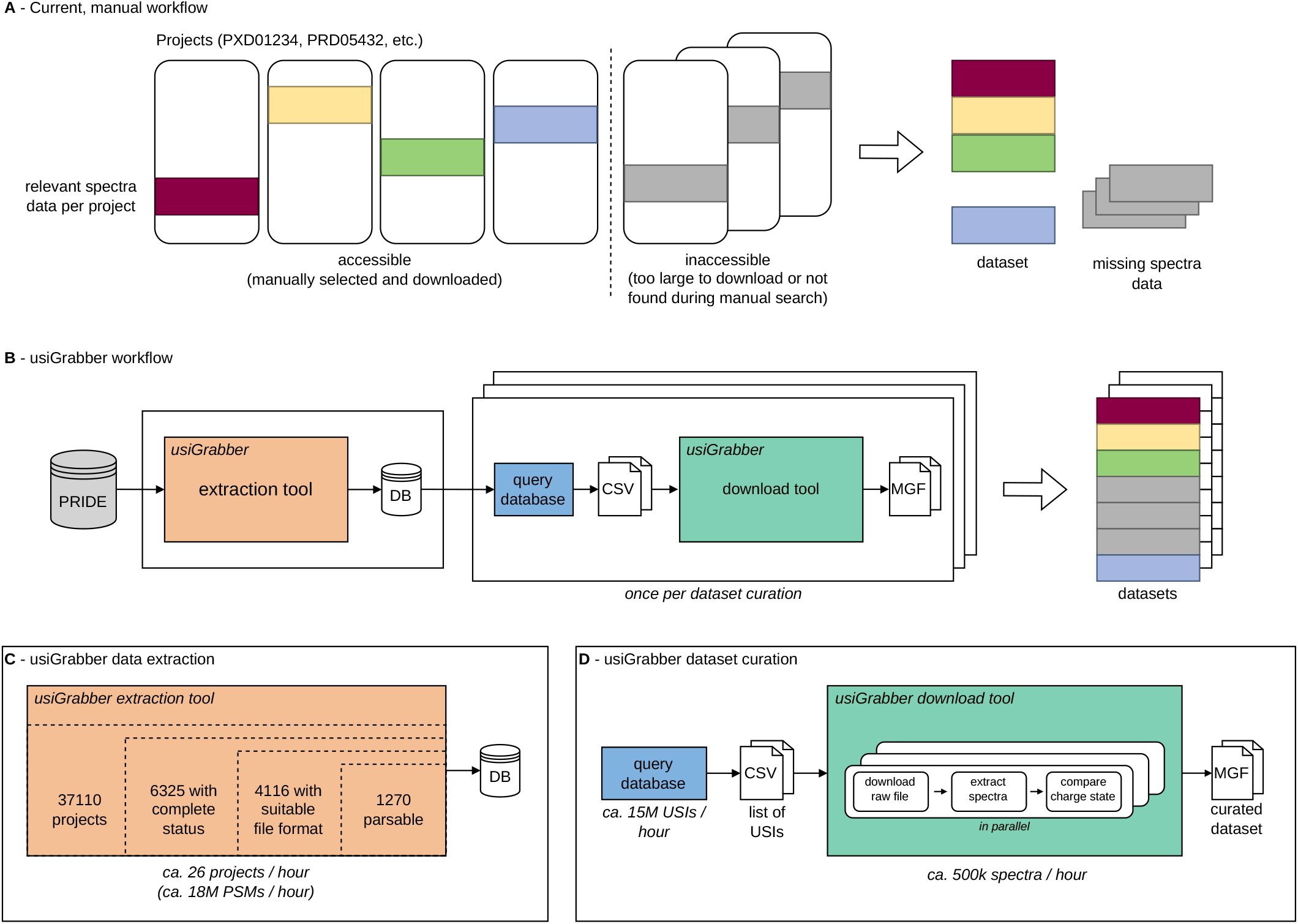
**A** outlines the current mass-spectrometry proteomics dataset curation. Various projects are downloaded in full, only filtered by project-level metadata. From the projects accessible, research-relevant spectra are then combined into the final dataset. However, some projects may be excluded if they are too large or cannot be found, even if their spectra would otherwise meet the criteria. **B** provides an overview of the usiGrabber workflow. The extraction tool creates a database to allow in-depth filtering of spectra data available on PRIDE. This allows filtering for research-relevant USIs, for which the download tool provides the corresponding spectra. Thus, this workflow creates a dataset containing all relevant spectra from all projects in the usiGrabber database, including spectra data that would not have been discovered during manual data curation. Also, once the extraction tool has built the database, various datasets can be curated by different queries. **C** shows how the usiGrabber extraction tool narrows down the PRIDE database to 1,270 projects that follow our data quality standards. With an approximate extraction speed of 18 million PSMs per hour, our tool parsed these projects within 2 days and populated the usiGrabber database with over 800 million PSMs. **D** The dataset curation process queries our database for a suitable list of USIs, which the download tool subsequently uses to acquire corresponding spectra. The download tool retrieves individual raw files from PRIDE and extracts all relevant spectra, which undergo a final quality control step to ensure their validity. This process leverages massive parallelization to guarantee maximum efficiency. The dataset curation of our proof-of-concept dataset showed download speeds of around 500,000 spectra per hour, creating a machine learning-ready dataset in about 30 hours. Overall, the usiGrabber allows for a significantly faster and easier curation of proteomic datasets compared to current manual workflows. Additionally, our workflow ensures that all projects available in the database are considered for the dataset curation, so we do not miss relevant spectra data.

A key reason for this is that current public repositories are optimized for deposition and project-level discovery rather than for spectrum-level retrieval and dataset assembly. PRIDE centrally hosts a large fraction of public proteomics data, but its search and filtering options are largely limited to coarse project metadata such as organism, instrument, or keywords (14). For many downstream tasks, the relevant information is either distributed across result files or only becomes apparent at the spectrum or Peptide Spectrum Match (PSM) level (e.g., confident site-localized modifications, charge states, fragmentation modes, or per-spectrum quality indicators). Even after manually selecting projects, building a high-quality combined dataset remains challenging due to heterogeneity in instruments, acquisition strategies, search parameters, and confidence thresholds.

Mass Spectromy Interactive Virtual Environment (MassIVE) (3) provides a complementary ecosystem and goes one step further with reanalysis pipelines to harmonize identification and quality control across submissions. With MassIVE-KB (22), it offers a widely used dataset of reanalysed spectra that has enabled multiple downstream models. Yet, this approach comes with trade-offs that limit its accessibility and flexibility for rapid dataset prototyping. Reanalysis is inherently storage- and compute-intensive and, therefore, realistically available to only a small number of groups. Moreover, reanalysis-centric pipelines are best suited when a full reprocessing of entire projects is required, but they become inefficient when only subsets of projects or files are relevant. The publicly exposed MassIVE selection mechanisms provide only basic filtering, which makes it difficult to extract narrowly defined subsets without consuming very large fractions of the corpus. Limited service availability prevented us from successfully downloading the complete MassIVE-KB V2 release. Furthermore, attempting to download larger specific subsets, such as phosphorylations, resulted in database crashes.

In a unification effort, the Universal Spectrum Identifier (USI) standard (4) was introduced to make individual spectra citable and referenceable across repositories. However, the existence of a spectrum identifier alone does not solve the practical problem of retrieving spectra at scale and turning them into machine learning-ready datasets: Tools like pridepy (7) only provide bulk download of entire raw files, while the PRIDE Archive USI service (15) is solely intended for the visualization of individual USIs. As a result, data-preparation workflows still require numerous manual steps to convert a set of interesting projects into a consistent dataset with labels and metadata.

We propose **usiGrabber**, a scalable framework for assembling large and diverse proteomics machine learning datasets that is designed around portability and extensibility rather than around a single centrally maintained database. The core idea is a light-weight *result-file parser* approach: Instead of requiring full reanalysis, usiGrabber extracts spectrum-level evidence and metadata from existing repository artifacts (e.g., identification result files), normalizes them into a common schema, and indexes them by USIs. This enables a workflow where researchers can (i) construct their own database instance from repositories and result files they care about, (ii) filter candidates at the spectrum/PSM level using rich metadata, and (iii) materialize the matched USIs into training-ready datasets in a scalable way. Importantly, our goal is not to release a single fixed dataset, but to provide reusable and extensible software that enables others to construct task-specific datasets on commodity infrastructure.

We implement a robust and extensible mzIdentML parser and use usiGrabber to construct a task-specific dataset targeting the presence of phosphorylations. We then retrain a spectrum classification model on this dataset and achieve comparable performance to the published baselines. Notably, the end-to-end process demonstrates a practical, scalable workflow for rapidly prototyping and benchmarking task-specific machine learning models in proteomics—with the ability to incorporate newly published data as it becomes available, rather than being tied to a small set of legacy datasets.

## Methods

**usiGrabber** automates the curation of proteomics spectra data at scale, making large datasets ready for use in machine learning systems. It consists of three main steps, as shown in Figure 1B:

1. Extracting metadata and PSMs once from projects on PRIDE.
2. Filtering the extracted data and identifying relevant spectrum identifiers for each specific research question, respectively.
3. Downloading spectra at scale based on the filtered USIs.

To demonstrate this functionality, we curate a dataset for training a binary phosphorylation classifier.

### Data sources

We focus on data from PRIDE since it is the leading repository for spectra data (15) and offers a well-documented API.

Project-level metadata is extracted directly from the PRIDE API, but detailed spectrum identification data is stored in either *search* files or *result* files. Search files include tool-specific formats from Mascot (16), MaxQuant (19), and more, whereas result files use standardized formats from the HUPO Proteomics Standards Initiative: *mzIdentML* (2) and *mzTab* (5). The mzTab format provides a simpler, but less complete overview compared to mzIdentML files, which is why we focus on mzIdentML files only. Projects that contain result files are considered complete and ensure an intact link between spectrum identification data and the spectrum itself (18). As visualized in Figure 1C, 6,325 out of the 37,110 currently accessible projects are such complete projects. To simplify retrieving raw spectra, we only consider projects providing Thermo Fisher Raw files, which are available in over 95 % of projects in PRIDE. After restricting complete projects to those providing both mzIdentML and raw files, we end up with 4,116 projects in total.

### Extracting spectrum identification data and project metadata

For each project, we download and parse mzIdentML files and store a relevant subset of the available information. Specifically, we extract file-level metadata, such as the analysis software and quality-control thresholds used. For each PSM, we further extract the following:

- Name of the corresponding raw file and the associated scan number, which are required to construct the USI
- Matched peptide sequence and post-translational modifications
- Identified proteins that include the peptide
- Various other information, such as charge state, calculated mass-to-charge ratio, and experimental mass-to-charge ratio

Our parser extends mzIdentML parsing utilities from *pyteomics* (10) with robust error handling and additional adaptive parsing for various cases in which the exact formatting deviates from the standard. Still, the extraction of the relevant fields for spectrum retrieval is a complex task. As file standards and tool versions change over time, so does the data format. While these variations may differ in their extent, it is important to take these into account when creating parsers or adapting existing ones for future data format changes. Project-level metadata, such as project accession, organisms, and instruments, do not require additional parsing capabilities and can be extracted directly from the PRIDE API.

For both spectrum-level and project-level data, we leverage ontologies such as UNIMOD, NCBITaxon, and MS through the *EMBL-EBI Ontology Lookup Service* (12) to enrich and unify these annotations. Additionally, we use identifiers from the *Universal Protein Knowledgebase (UniProt)* (20) to link protein information.

The overall setup is flexible and includes tooling for retrying failed file or project extraction attempts, as well as a mechanism to easily add future projects.

### Querying USIs and downloading spectra at scale

With the usiGrabber database in place, numerous datasets can be created by filtering the extracted data for relevant PSMs and associated USIs, which is done with simple queries. However, existing tools for retrieving raw spectra based on USIs like the PRIDE USI service (15) are optimized for visualization of individual spectra and do not scale in practice. The usiGrabber download tool, visualized in Figure 1D, closes this gap by grouping USIs by raw file and downloading raw files from PRIDE. Spectra are then extracted with version 2.0 of the ThermoRawFileParser (6) and further post-processed to produce Parquet or MGF files.

During this process, we observed minor inconsistencies between USIs built from information in mzIdentML files and the raw spectra, necessitating a quality control step. Although raw spectra contain limited additional metadata, the charge state is available in both the spectrum and the USI. We therefore compare both values to validate that linking is correct and discard all raw files with at least one charge-state mismatch. Since there is no certain way to know if a single faulty spectrum is an isolated anomaly or an indicator of a larger data quality issue, this is the only way to ensure maximum confidence in the correctness of all downloaded spectra.

### Training a binary phosphorylation classifier

As a proof of concept, we use the usiGrabber to curate a dataset and train a binary phosphorylation classifier based on the AHLF model architecture proposed by Altenburg et al. (1). It is a deep learning model that is able to detect the presence of a phosphorylation from raw spectrum data.

To construct a dataset, we filter the usiGrabber database for sequences with and without detected phosphorylations from experiments in which a search for phosphorylations was performed. In addition to our quality control step, we filter for USIs whose corresponding PSM passes the project-specific quality threshold and retain only the best-matching search results (rank equal to 1).

For each raw file, we create one corresponding Parquet and MGF file containing spectra data. To avoid potential ordering biases in the data, we apply seeded bucket-based shuffling as a postprocessing step and obtain roughly randomly selected 5,000 PSMs in each file. During training, we apply a 93/7 train/validation split.

We use *MassIVE-KB 2*.*0*.*15, in vivo, Modified only* (23) as our independent test set. For both our dataset and the test set, we verify that there is no overlap in individual USIs, MS run names, and project accessions. Thus, we ensure that our evaluation is not misguided by any potential data quality issues or biases introduced by our process.

All training runs based on the AHLF model architecture ran for 10 virtual epochs with a batch size of 128 and 37, 000 steps each. Batches were balanced to contain equal numbers of phosphorylated and non-phosphorylated PSMs. The hyperparameters optimized were the learning rate and the ion intensity normalization function. Normalizing by the maximum intensity consistently yielded faster convergence and lower validation loss on our data than the squared-sum normalization used in the original AHLF. The final model is selected by maximizing the harmonic mean of the accuracy for both class validation scores on a 0.99 time weighted average.

### Compute and storage infrastructure

The usiGrabber extraction tool is deployed on a server with 16 CPU cores, 265 GB RAM, 500 GB storage, and a 25 Gbps network connection. Extracted data is stored in a standard PostgreSQL database hosted on a second server with 16 CPU cores, 32 GB RAM, and 2 TB of SSD storage. Querying the database for relevant USIs and downloading spectra requires no special infrastructure, but benefits from a fast network connection. To adhere to fair usage best practices and to not overuse the PRIDE infrastructure, we intentionally limit download speeds for both data extraction and spectra retrieval. For training the binary phosphorylation classifier, we used a single V100 GPU.

## Results

The execution of the usiGrabber extraction tool took roughly 48 hours. It resulted in a database, encompassing more than 800 million PSMs across more than 1,200 different projects. On average, we have a throughput of 26 PRIDE projects or 18 million PSMs per hour. As shown in Figure 2, the database comprises a wide variety of data. Figure 2A shows the diversity of sampled species. Ontology information can be used to further enhance the filtering options, which allows filtering for broader search terms, such as bacteria. Similar to sample species, the database spans a variety of instrument manufacturers and types associated with spectra data, as shown in Figure 2B.

**Figure 2.**
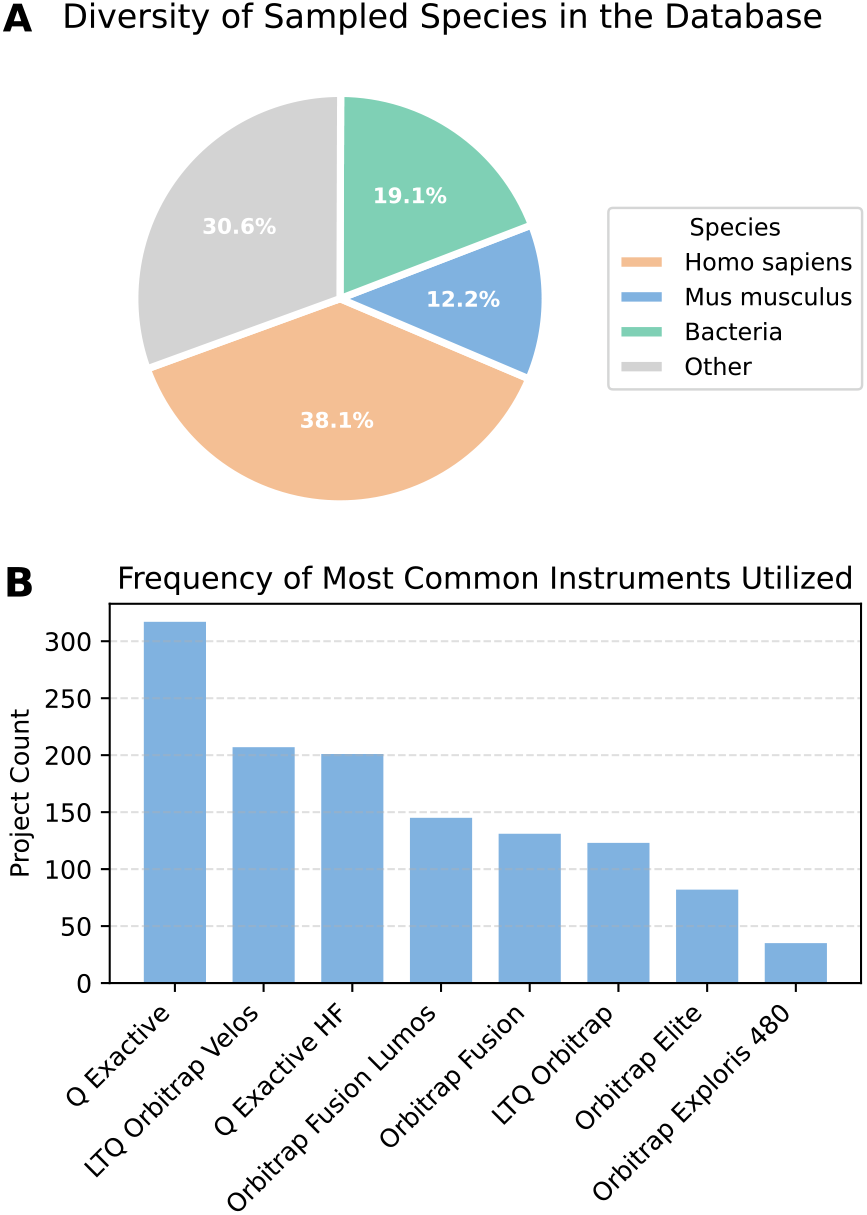
**A** illustrates the distribution of projects according to the species sampled across all 1,270 distinct projects in our database. B displays the frequency of the most commonly used instruments over the same projects. Together, these figures underscore the diversity and practical scope of data captured by usiGrabber.

Although the PRIDE database contains around 6, 000 complete projects, not all of them allow for straightforward identification and retrieval of relevant spectra. Thus, we focus only on projects that include mzIdentML data as well as spectra data in *Thermo Fisher Raw* format and are parseable. Figure 3A shows the number of complete projects in PRIDE, which contain at least one mzIdentML file, in contrast to those that contain none. After an initial drop in projects that provide their results exclusively in mzTab format, there has been another rise in mzTab-only projects recently (see Figure 3B). In an effort to capture the largest volume of available data, the usiGrabber currently focuses on projects containing mzIdentML files, which leaves us with 4, 116 suitable projects. Out of these, our extraction tool successfully parsed and validated 1, 270 projects.

**Figure 3.**
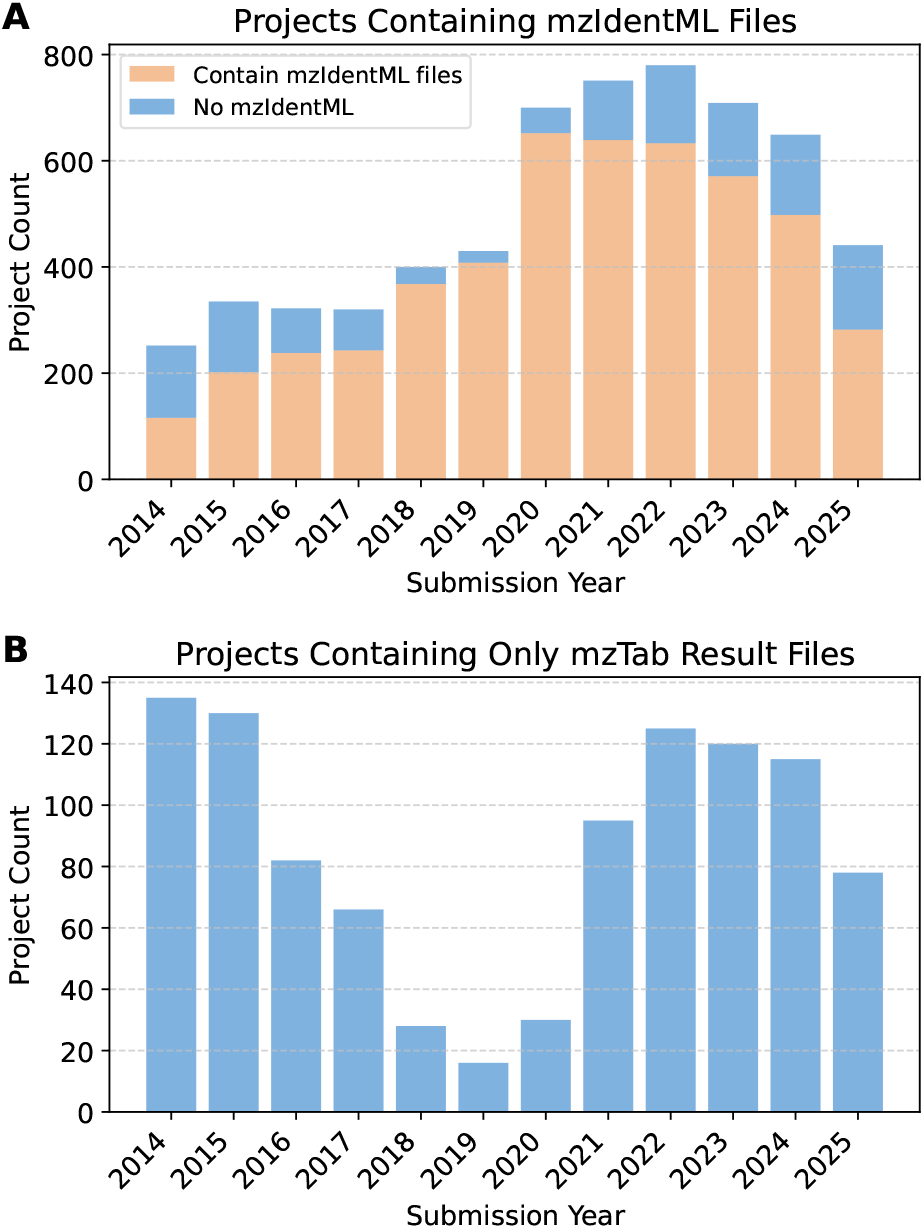
**A** shows the number of projects in PRIDE with complete status per year and how many of them contain mzIdentML files. **B** visualizes solely the amount of those projects that do not contain mzIdentML, but rather only mzTab files. As only mzIdentML files hold all the information for USI reconstruction, we implemented processing exclusively for them. However, since the number of complete projects only providing mzTab files has grown significantly since 2019, this has contributed to a lower count of successfully parsed projects in recent years.

### Binary phosphorylation classifier

We relied on the usiGrabber framework to curate a dataset for our task-specific use case. To gather all relevant PSMs with and without phosphorylation for the classification task, we executed two distinct dataset-construction queries. In about 20 minutes, these queries yielded approximately 12 million PSMs with corresponding USIs. Downloading finished autonomously in roughly one day, and our dataset postprocessing ran for another 5 hours. All in all, our dataset was ready for machine learning in less than two working days with minimal human interaction. Our final dataset comprises 10, 961, 094 PSMs, narrowed from 42, 592, 824 potentially available PSMs (see Table 1). Due to our strict charge state consistency verification, approximately 57 % of PSMs are excluded in the final step. Although this conservative strategy likely excludes valid spectra and could be refined in future iterations, we deliberately prioritized robustness and correctness over maximum data utilization.

**Table 1.** Cumulative stepwise data quality control filters on phosphorylated and non-phosphorylated PSMs reduces 42,592,824 potentially available PSMs to 10,961,094 high-confidence PSMs used as the dataset for our model training.

| Filters | Phosphorylation | Other |
| --- | --- | --- |
| None | 18,753,926 | 23,838,898 |
| Rank is 1 | 17,029,070 | 22,894,086 |
| Passed threshold | 8,488,851 | 16,879,595 |
| Verified charge states | 4,912,911 | 6,048,183 |

Note: Each filter also includes all previous filters.

The original AHLF model was trained using a 4-fold cross-validation on nearly 20 million manually curated spectra and combined into one ensemble model for inference. Table 2 shows the test set performance of our final model compared to all AHLF models (Alpha, Beta, Delta, Gamma). With a balanced accuracy of 0.78, our model performs on a comparable performance basis with all individual AHLF models. We minimally outperform the mean of the balanced accuracies of the AHLF models and have less than 1 % difference to the median balanced accuracy. Only the combined ensemble model, which uses twice as much training data, beats our model by approximately 3 percentage points on balanced accuracy and F1 score.

**Table 2.** Binary Phosphorylation Classification Model Performance Comparison between AHLF (as published) as reference and our model. The comparison is based on the balanced accuracy and F1 score and shows that our model performs comparably to AHLF Combined, beating two of four models (Alpha and Delta) on both metrics.

| Category | Model | Balanced Accuracy | F1 Score |
| --- | --- | --- | --- |
| Reference (AHLF) | AHLF Combined | **0.8094** | **0.7421** |
|  | AHLF Alpha | 0.7543 | 0.6802 |
|  | AHLF Beta | 0.7933 | 0.7276 |
|  | AHLF Delta | 0.7813 | 0.7095 |
|  | AHLF Gamma | 0.8050 | 0.7364 |
| Our Model | usiGrabber AHLF | 0.7822 | 0.7114 |

## Discussion

In this work, we introduce **usiGrabber**, a scalable framework designed to bridge the gap between vast, often inaccessible public proteomics repositories and the specialized requirements of modern machine learning. Mass spectrometry-based proteomics is uniquely positioned to provide foundational biological insights into cellular status, offering details on post-translational modification (PTM) landscapes that are largely invisible to genomic or transcriptomic layers. As demonstrated by recent high-resolution mapping of disease-specific PTMs (9), capturing these functional states is essential for characterizing complex biological systems and their current phenotypic manifestations.

Despite this importance, MS-based proteomics has yet to fully benefit from the transformative shift toward artificial intelligence that has redefined genomics. While Protein Language Models (PLMs) have fundamentally changed the prediction of protein structure and design from sequences (11), their application to raw proteomics data remains constrained by a significant data access bottleneck. Although the community has spent two decades establishing open data sharing, the resulting repositories remain heterogeneous and difficult to compile for targeted ML tasks. **usiGrabber** addresses this by enabling a shift from static, decade-old legacy training sets toward dynamic, automated extraction. We demonstrate that task-specific datasets containing millions of spectra can be curated in under two days, offering a scalable alternative to resource-intensive bulk download strategies or tedious manual curation. We demonstrate that these advantages in usability do not negatively impact data quality.

While the current reliance on `mzIdentML` formats and the inherent noise of metadata present ongoing hurdles, **usiGrabber** provides the modular infrastructure needed to overcome these technical bottlenecks. By integrating emerging metadata standards like Sample and Data Relationship Format (SDRF) (21) and expanding support for search-specific file types like MaxQuant, the framework is poised to scale alongside the rapid growth of public repositories. Ultimately, by replacing limited synthetic data with large-scale, real-world MS datasets, this work provides a necessary link for proteomics-based ML to reach a scale comparable to its sequence-based counterparts.

## Acknowledgments

We thank participants of an internal hackathon: Jakub M. Bartoszewicz, Katharina Baum, Andrea Eoli, Henrike O. Heyne, Pauline Hiort, Pascal Iversen, Kunaphas Kongkitimanon, Marta S. Lemanczyk, Fábio Miranda, Ferdous Nasri, Melania Nowicka, Pia Francesca Rissom, Jan-Philipp Sachs, Anna-Juliane Schmachtenberg, Linea Schmidt, Jens-Uwe Ulrich, Julian S. Wanner, Simon Witzke, Paulo Yanez, Elizabeth Y. Yuu, and Justus Zeinert.

This work is supported by a European Research Council (ERC) grant (eXplAInProt, 101124385) to BYR.

## Author contributions

Georg Auge, Matthis Clausen, Konstantin Ketterer, Jacob Schaefer and Nils Schmitt (Data curation, Investigation, Formal analysis, Software, Visualization, Writing – original draft, Writing – review & editing), Tom Altenburg (Conceptualization), Yannick Hartmaring and Hendrik Raetz (Conceptualization, Supervision, Writing – review & editing), Bernhard Y. Renard and Christoph N. Schlaffner (Project administration, Conceptualization, Supervision, Writing – review & editing)

## Data and code availability

The code for the usiGrabber, as well as the model weights of our trained classifier, are available on GitHub (https://github.com/usiGrabber/usiGrabber). The repository also includes instructions on how to set up the database in its current state, which was exported and uploaded to https://zenodo.org/records/18853258. The dataset we curated for training the binary phosphorylation classifier is also available at https://zenodo.org/records/18853258. A full list of the used PRIDE project IDs is available in the supplementary material.

## Footnotes

1 https://www.ebi.ac.uk/pride/ws/archive/v3/stats/submitted-data

2 https://pubmed.ncbi.nlm.nih.gov/?term=machine+learning+proteomics

