## Supplemental material for "usiGrabber: Automating the curation of proteomics spectra data at scale, making large datasets ready for use in machine learning systems"

### Supplementary material

We extracted data from the following 1,270 projects using the usiGrabber extraction tool: PXD000164, PXD000329, PXD000402, PXD000498, PXD000710, PXD000780, PXD000791, PXD000854, PXD000897, PXD000966, PXD000968, PXD000997, PXD001000, PXD001002, PXD001006, PXD001057, PXD001072, PXD001172, PXD001177, PXD001179, PXD001212, PXD001213, PXD001267, PXD001340, PXD001344, PXD001357, PXD001362, PXD001447, PXD001482, PXD001483, PXD001509, PXD001511, PXD001518, PXD001519, PXD001520, PXD001568, PXD001572, PXD001578, PXD001714, PXD001726, PXD001827, PXD001830, PXD001884, PXD001885, PXD001924, PXD001926, PXD001928, PXD001951, PXD001968, PXD001974, PXD002021, PXD002026, PXD002040, PXD002041, PXD002042, PXD002043, PXD002044, PXD002045, PXD002046, PXD002047, PXD002048, PXD002049, PXD002050, PXD002080, PXD002081, PXD002082, PXD002083, PXD002084, PXD002085, PXD002086, PXD002087, PXD002088, PXD002089, PXD002161, PXD002163, PXD002193, PXD002253, PXD002266, PXD002308, PXD002309, PXD002310, PXD002311, PXD002312, PXD002317, PXD002346, PXD002365, PXD002372, PXD002440, PXD002470, PXD002471, PXD002486, PXD002538, PXD002588, PXD002633, PXD002655, PXD002658, PXD002679, PXD002681, PXD002694, PXD002763, PXD002772, PXD002781, PXD002788, PXD002789, PXD002796, PXD002798, PXD002841, PXD002855, PXD002883, PXD002901, PXD002944, PXD002982, PXD002986, PXD002990, PXD003022, PXD003023, PXD003029, PXD003037, PXD003045, PXD003055, PXD003079, PXD003182, PXD003200, PXD003232, PXD003346, PXD003356, PXD003599, PXD003639, PXD003666, PXD003672, PXD003676, PXD003731, PXD003826, PXD003827, PXD003828, PXD003829, PXD003830, PXD003876, PXD003881, PXD003916, PXD003922, PXD003927, PXD003943, PXD003976, PXD004002, PXD004005, PXD004062, PXD004064, PXD004065, PXD004135, PXD004139, PXD004204, PXD004333, PXD004413, PXD004510, PXD004540, PXD004569, PXD004583, PXD004636, PXD004651, PXD004682, PXD004684, PXD004741, PXD004760, PXD004763, PXD004785, PXD004865, PXD004883, PXD004890, PXD004896, PXD004935, PXD005055, PXD005101, PXD005117, PXD005152, PXD005292, PXD005295, PXD005306, PXD005335, PXD005341, PXD005359, PXD005407, PXD005415, PXD005434, PXD005440, PXD005590, PXD005620, PXD005632, PXD005696, PXD005697, PXD005723, PXD005728, PXD005755, PXD005759, PXD005794, PXD005851, PXD005908, PXD005923, PXD005979, PXD005980, PXD006036, PXD006066, PXD006084, PXD006096, PXD006169, PXD006204, PXD006205, PXD006216, PXD006265, PXD006268, PXD006311, PXD006316, PXD006320, PXD006413, PXD006428, PXD006440, PXD006466, PXD006467, PXD006536, PXD006537, PXD006569, PXD006615, PXD006715, PXD006756, PXD006757, PXD006847, PXD006848, PXD006863, PXD006910, PXD006920, PXD006938, PXD006994, PXD007050, PXD007086, PXD007137, PXD007149, PXD007153, PXD007188, PXD007226, PXD007267, PXD007268, PXD007287, PXD007528, PXD007575, PXD007648, PXD007681, PXD007688, PXD007700, PXD007713, PXD007748, PXD007782, PXD007788, PXD007790, PXD007791, PXD007864, PXD007890, PXD008046, PXD008103, PXD008128, PXD008136, PXD008199, PXD008205, PXD008217, PXD008238, PXD008272, PXD008280, PXD008310, PXD008450, PXD008490, PXD008544, PXD008595, PXD008622, PXD008645, PXD008647, PXD008667, PXD008675, PXD008723, PXD008786, PXD008796, PXD008808, PXD008832, PXD008836, PXD008844, PXD008851, PXD008895, PXD008909, PXD008920, PXD008952, PXD009019, PXD009021, PXD009029, PXD009066, PXD009072, PXD009080, PXD009096, PXD009105, PXD009216, PXD009387, PXD009401, PXD009435, PXD009480, PXD009483, PXD009489, PXD009549, PXD009611, PXD009614, PXD009640, PXD009649, PXD009665, PXD009681, PXD009682, PXD009698, PXD009712, PXD009713, PXD009736, PXD009810, PXD009817, PXD009818, PXD009837, PXD009852, PXD009863, PXD009950, PXD009982, PXD010000, PXD010001, PXD010042, PXD010074, PXD010142, PXD010145, PXD010150, PXD010193, PXD010252, PXD010258, PXD010271, PXD010357, PXD010362, PXD010429, PXD010430, PXD010431, PXD010456, PXD010487, PXD010515, PXD010570, PXD010575, PXD010589, PXD010610, PXD010613, PXD010659, PXD010663, PXD010699, PXD010700, PXD010794, PXD010806, PXD010816, PXD010827, PXD010908, PXD010911, PXD010986, PXD010995, PXD011029, PXD011070, PXD011073, PXD011081, PXD011123, PXD011141, PXD011156, PXD011160, PXD011170, PXD011183, PXD011280, PXD011287, PXD011365, PXD011427, PXD011431, PXD011519, PXD011520, PXD011521, PXD011582, PXD011583, PXD011692, PXD011697, PXD011712, PXD011714, PXD011721, PXD011817, PXD011948, PXD011988, PXD011992, PXD012062, PXD012170, PXD012177, PXD012178, PXD012179, PXD012190, PXD012191, PXD012200, PXD012225, PXD012232, PXD012260, PXD012407, PXD012463, PXD012484, PXD012490, PXD012491, PXD012494, PXD012522, PXD012523, PXD012611, PXD012739, PXD012748, PXD012753, PXD012754, PXD012770, PXD012796, PXD012798, PXD012810, PXD012813, PXD012865, PXD012961, PXD012963, PXD012965, PXD012967, PXD012969, PXD012979, PXD012982, PXD012986, PXD012991, PXD013004, PXD013036, PXD013080, PXD013090, PXD013093, PXD013159, PXD013196, PXD013198, PXD013211, PXD013213, PXD013214, PXD013250, PXD013264, PXD013273, PXD013274, PXD013280, PXD013281, PXD013282, PXD013300, PXD013320, PXD013321, PXD013322, PXD013329, PXD013343, PXD013350, PXD013369, PXD013406, PXD013449, PXD013503, PXD013514, PXD013541, PXD013608, PXD013656, PXD013684, PXD013710, PXD013711, PXD013712, PXD013724, PXD013759, PXD013805, PXD013806, PXD013890, PXD013896, PXD013897, PXD013899, PXD013962, PXD013964, PXD014071, PXD014166, PXD014230, PXD014254, PXD014300, PXD014317, PXD014419, PXD014505, PXD014550, PXD014552, PXD014607, PXD014617, PXD014730, PXD014780, PXD014820, PXD014933, PXD014976, PXD015002, PXD015015, PXD015022, PXD015057, PXD015058, PXD015059, PXD015060, PXD015061, PXD015111, PXD015149, PXD015153, PXD015157, PXD015168, PXD015203, PXD015253, PXD015337, PXD015371, PXD015407, PXD015460, PXD015462, PXD015485, PXD015635, PXD015673, PXD015698, PXD015700, PXD015727, PXD015817, PXD015836, PXD015982, PXD016017, PXD016160, PXD016172, PXD016224, PXD016225, PXD016248, PXD016254, PXD016332, PXD016377, PXD016394, PXD016442, PXD016446, PXD016456, PXD016487, PXD016557, PXD016560, PXD016629, PXD016648, PXD016697, PXD016698, PXD016876, PXD016930, PXD016964, PXD016987, PXD017017, PXD017063, PXD017115, PXD017119, PXD017148, PXD017150, PXD017198, PXD017229, PXD017239, PXD017260, PXD017271, PXD017308, PXD017392, PXD017469, PXD017518, PXD017535, PXD017596, PXD017650, PXD017657, PXD017659, PXD017677, PXD017709, PXD017747, PXD017792, PXD017847, PXD017891, PXD017943, PXD017964, PXD017993,

PXD018005, PXD018045, PXD018061, PXD018067, PXD018079, PXD018123, PXD018322, PXD018329, PXD018372, PXD018374, PXD018517, PXD018578, PXD018602, PXD018644, PXD018672, PXD018701, PXD018749, PXD018826, PXD018897, PXD018945, PXD018952, PXD018953, PXD018954, PXD018955, PXD018968, PXD018981, PXD018987, PXD019015, PXD019058, PXD019093, PXD019095, PXD019149, PXD019219, PXD019238, PXD019317, PXD019327, PXD019335, PXD019410, PXD019464, PXD019472, PXD019474, PXD019491, PXD019541, PXD019559, PXD019605, PXD019610, PXD019622, PXD019676, PXD019686, PXD019693, PXD019771, PXD019824, PXD019825, PXD019830, PXD019831, PXD019839, PXD019846, PXD019850, PXD019851, PXD019897, PXD019898, PXD019908, PXD019911, PXD019912, PXD019913, PXD019928, PXD019931, PXD019952, PXD020100, PXD020124, PXD020152, PXD020175, PXD020186, PXD020210, PXD020219, PXD020241, PXD020250, PXD020269, PXD020366, PXD020456, PXD020463, PXD020499, PXD020516, PXD020520, PXD020547, PXD020593, PXD020608, PXD020677, PXD020737, PXD020767, PXD020778, PXD020868, PXD020976, PXD020998, PXD021069, PXD021123, PXD021163, PXD021181, PXD021194, PXD021201, PXD021221, PXD021227, PXD021233, PXD021271, PXD021306, PXD021332, PXD021368, PXD021422, PXD021431, PXD021498, PXD021516, PXD021580, PXD021606, PXD021661, PXD021676, PXD021759, PXD021816, PXD021817, PXD021837, PXD021855, PXD021897, PXD021974, PXD021992, PXD022049, PXD022074, PXD022106, PXD022163, PXD022196, PXD022244, PXD022284, PXD022287, PXD022300, PXD022353, PXD022377, PXD022407, PXD022555, PXD022570, PXD022676, PXD022767, PXD022770, PXD022824, PXD022858, PXD022953, PXD023034, PXD023043, PXD023096, PXD023178, PXD023262, PXD023265, PXD023354, PXD023359, PXD023459, PXD023514, PXD023516, PXD023548, PXD023626, PXD023692, PXD023702, PXD023810, PXD023812, PXD023913, PXD023922, PXD023974, PXD024020, PXD024059, PXD024065, PXD024116, PXD024138, PXD024159, PXD024201, PXD024482, PXD024483, PXD024484, PXD024485, PXD024582, PXD024583, PXD024584, PXD024620, PXD024627, PXD024634, PXD024638, PXD024688, PXD024702, PXD024708, PXD024714, PXD024723, PXD024791, PXD024796, PXD024840, PXD024846, PXD024847, PXD024848, PXD024849, PXD024850, PXD024888, PXD024927, PXD024968, PXD024990, PXD024992, PXD024993, PXD025004, PXD025008, PXD025030, PXD025130, PXD025131, PXD025141, PXD025210, PXD025235, PXD025238, PXD025264, PXD025471, PXD025504, PXD025505, PXD025641, PXD025808, PXD025809, PXD025888, PXD025967, PXD026063, PXD026116, PXD026189, PXD026431, PXD026461, PXD026506, PXD026513, PXD026618, PXD026651, PXD026725, PXD026793, PXD026798, PXD026833, PXD026875, PXD026895, PXD026953, PXD027034, PXD027076, PXD027105, PXD027122, PXD027158, PXD027327, PXD027351, PXD027453, PXD027491, PXD027618, PXD027706, PXD027728, PXD027794, PXD027796, PXD027803, PXD027823, PXD027827, PXD027867, PXD027923, PXD027944, PXD027955, PXD027964, PXD028074, PXD028161, PXD028272, PXD028322, PXD028379, PXD028495, PXD028514, PXD028538, PXD028576, PXD028584, PXD028595, PXD028596, PXD028639, PXD028685, PXD028802, PXD028827, PXD028833, PXD028840, PXD028846, PXD028867, PXD028917, PXD028933, PXD029006, PXD029267, PXD029289, PXD029424, PXD029428, PXD029432, PXD029453, PXD029490, PXD029577, PXD029783, PXD029850, PXD029901, PXD029959, PXD029979, PXD029998, PXD030077, PXD030114, PXD030118, PXD030161, PXD030163, PXD030165, PXD030170, PXD030293, PXD030319, PXD030331, PXD030467, PXD030468, PXD030545, PXD030610, PXD030684, PXD030748, PXD030859, PXD030861, PXD030888, PXD030889, PXD030900, PXD030923, PXD031004, PXD031012, PXD031022, PXD031046, PXD031065, PXD031072, PXD031166, PXD031261, PXD031271, PXD031346, PXD031359, PXD031361, PXD031379, PXD031457, PXD031515, PXD031524, PXD031570, PXD031583, PXD031656, PXD031683, PXD031766, PXD031772, PXD031786, PXD031802, PXD032203, PXD032295, PXD032356, PXD032387, PXD032800, PXD032844, PXD032920, PXD033040, PXD033105, PXD033175, PXD033180, PXD033330, PXD033354, PXD033387, PXD033408, PXD033486, PXD033494, PXD033519, PXD033536, PXD033581, PXD033603, PXD033607, PXD033691, PXD033692, PXD033887, PXD033912, PXD033941, PXD033943, PXD034035, PXD034089, PXD034117, PXD034138, PXD034159, PXD034161, PXD034179, PXD034365, PXD034376, PXD034494, PXD034577, PXD034578, PXD034579, PXD034587, PXD034601, PXD034602, PXD034628, PXD034795, PXD034825, PXD034840, PXD034846, PXD034903, PXD034932, PXD034938, PXD034980, PXD035019, PXD035020, PXD035172, PXD035201, PXD035255, PXD035305, PXD035526, PXD035569, PXD035707, PXD035728, PXD035743, PXD035767, PXD035849, PXD035863, PXD035870, PXD036038, PXD036079, PXD036182, PXD036266, PXD036267, PXD036295, PXD036325, PXD036392, PXD036436, PXD036636, PXD036716, PXD036742, PXD036771, PXD036772, PXD036789, PXD036822, PXD037036, PXD037093, PXD037117, PXD037132, PXD037293, PXD037474, PXD037546, PXD037575, PXD037674, PXD037894, PXD037920, PXD038009, PXD038010, PXD038065, PXD038068, PXD038128, PXD038147, PXD038237, PXD038248, PXD038303, PXD038319, PXD038327, PXD038492, PXD038494, PXD038549, PXD038554, PXD038636, PXD038644, PXD038703, PXD038712, PXD038727, PXD038742, PXD038746, PXD038849, PXD038898, PXD038934, PXD039007, PXD039053, PXD039467, PXD039499, PXD039577, PXD039652, PXD040094, PXD040102, PXD040121, PXD040185, PXD040187, PXD040223, PXD040344, PXD040345, PXD040499, PXD040565, PXD040650, PXD040803, PXD040943, PXD041053, PXD041094, PXD041156, PXD041327, PXD041331, PXD041415, PXD041463, PXD041471, PXD041472, PXD041495, PXD041690, PXD041780, PXD041786, PXD042059, PXD042074, PXD042374, PXD042384, PXD042474, PXD042545, PXD042682, PXD042729, PXD042735, PXD042904, PXD043218, PXD043240, PXD043403, PXD043460, PXD043653, PXD043814, PXD043920, PXD043940, PXD043979, PXD044034, PXD044078, PXD044079, PXD044118, PXD044149, PXD044187, PXD044196, PXD044234, PXD044269, PXD044270, PXD044303, PXD044625, PXD044755, PXD044759, PXD044913, PXD044927, PXD045023, PXD045274, PXD045312, PXD045316, PXD045331, PXD045395, PXD045451, PXD045556, PXD045611, PXD045647, PXD045663, PXD045670, PXD045803, PXD045824, PXD045838, PXD045873, PXD045922, PXD045957, PXD046001, PXD046098, PXD046217, PXD046271, PXD046359, PXD046708, PXD046729, PXD046824, PXD046830, PXD046892, PXD046895, PXD046898, PXD046911, PXD047139, PXD047334, PXD047533, PXD047572, PXD047617, PXD047711, PXD047715, PXD047724, PXD047824, PXD047971, PXD047996, PXD048015, PXD048215, PXD048319, PXD048360, PXD048362, PXD048367, PXD048568, PXD048765, PXD048810, PXD048927, PXD049324, PXD049349, PXD049359, PXD049416, PXD050108, PXD050212, PXD050244, PXD050318, PXD050420, PXD050483, PXD050601, PXD050817,

PXD051099, PXD051157, PXD051183, PXD051310, PXD051418, PXD051472, PXD051750, PXD051905, PXD052138, PXD052144, PXD052186, PXD052289, PXD052317, PXD052442, PXD052621, PXD052639, PXD052694, PXD052797, PXD052839, PXD052869, PXD053040, PXD053068, PXD053100, PXD053107, PXD053108, PXD053134, PXD053439, PXD053444, PXD053505, PXD053593, PXD054011, PXD054267, PXD054269, PXD054272, PXD054278, PXD054280, PXD054282, PXD054283, PXD054285, PXD054317, PXD054418, PXD054475, PXD054716, PXD054747, PXD055080, PXD055133, PXD055167, PXD055236, PXD055244, PXD055293, PXD055392, PXD055393, PXD055411, PXD055768, PXD055777, PXD055830, PXD055838, PXD056020, PXD056179, PXD056446, PXD056732, PXD056752, PXD056779, PXD056910, PXD057140, PXD057691, PXD058079, PXD058419, PXD058429, PXD058865, PXD058911, PXD059520, PXD059538, PXD060475, PXD060651, PXD060734, PXD060738, PXD061013, PXD061097, PXD061212, PXD061237, PXD061267, PXD061282, PXD061530, PXD061593, PXD061651, PXD061836, PXD061882, PXD062120, PXD062128, PXD062210, PXD063113, PXD063191, PXD063192, PXD063569, PXD063761, PXD064480, PXD064940, PXD067360, PXD067854, PXD068145, PXD068189, PXD068228, PXD068413, PXD069333, PXD069609, and PXD071040.

We used data from the following 114 projects for retraining the binary phosphorylation classifier: PXD000997, PXD001000, PXD001057, PXD001072, PXD001177, PXD001179, PXD001340, PXD001344, PXD001447, PXD001572, PXD001578, PXD001830, PXD002266, PXD002471, PXD002486, PXD002538, PXD002796, PXD002990, PXD003232, PXD003599, PXD003922, PXD004636, PXD005152, PXD005359, PXD005415, PXD006036, PXD006268, PXD006413, PXD007050, PXD007137, PXD008832, PXD008952, PXD009401, PXD009698, PXD009863, PXD010145, PXD010362, PXD010570, PXD010589, PXD010827, PXD011156, PXD011170, PXD011431, PXD011583, PXD012191, PXD012407, PXD012753, PXD012754, PXD012770, PXD012813, PXD012963, PXD013503, PXD013964, PXD014552, PXD014976, PXD015059, PXD015060, PXD015635, PXD015673, PXD015836, PXD017119, PXD017308, PXD017469, PXD017847, PXD017943, PXD018374, PXD018672, PXD018981, PXD018987, PXD019093, PXD020210, PXD020241, PXD020250, PXD021368, PXD021992, PXD022106, PXD022676, PXD023096, PXD023974, PXD024846, PXD025878, PXD025888, PXD026461, PXD026618, PXD027034, PXD028595, PXD028827, PXD028840, PXD028917, PXD030467, PXD031515, PXD031570, PXD032920, PXD033912, PXD034089, PXD035526, PXD035569, PXD035707, PXD036325, PXD036742, PXD037117, PXD037132, PXD038934, PXD045556, PXD046281, PXD046729, PXD046824, PXD046830, PXD047996, PXD048215, PXD048765, PXD050212, PXD055411, and PXD056821.

Note that the projects PXD025878, PXD046281, and PXD056821 are not included above. This is due to the fact that we created the final usiGrabber extraction database after curating a dataset for the binary phosphorylation classifier.
